# A lipid cue drives the subcellular localization of a self-inserting bacterial transmembrane protein

**DOI:** 10.64898/2026.05.27.728190

**Authors:** Vani Pande, Taylor B. Updegrove, Vivek Anantharaman, Ashley Bae, Jiji Chen, L. Aravind, Kumaran S. Ramamurthi

## Abstract

The correct subcellular localization of proteins is a critical step underlying myriad biological processes, but the cues that drive the specific localization of integral membrane proteins in bacteria remain largely undeciphered. During sporulation in *Bacillus subtilis*, a rod-shaped outer “mother cell” constructs an internal spherical “forespore” cell that eventually matures into the spore. The integral membrane protein ShfA is made in the mother cell cytosol and localizes to the surface of the forespore. Here, we report that, despite being a multi-pass transmembrane protein, ShfA spontaneously inserts into the lipid bilayer via its N-terminal “YabQ” domain without the apparent need for a pre-localized insertase. ShfA preferentially inserts into cell division septa in multiple bacterial species, indicating that a widely conserved septal cue drives ShfA localization. Structural modeling suggested that the YabQ domain harbors a specific intramembrane groove that can bind the universal lipid carrier undecaprenyl phosphate (UndP), and UndP depletion in vivo disrupted proper ShfA localization. We propose that ShfA localizes to sites of active cell wall synthesis by binding to UndP and/or molecules like lipid I and lipid II that contain UndP and speculate that the function of ShfA is to stabilize these precursors of cell wall biogenesis from the harsh cytosolic nanoenvironment that surrounds the forespore during sporulation.

## INTRODUCTION

Despite their diminutive size and typical lack of membrane-bound organelles, bacterial cells exhibit high levels of internal organization, which may manifest as the characteristic subcellular localization patterns of specific proteins (1, 2). In bacteria, the proper localization of proteins in the cytosol underlies multiple cellular processes, such as the assembly of larger protein complexes that drive cell division, growth, differentiation, and morphogenesis, and the relocation of proteins in response to environmental signals (3–12). In the absence of a formal protein trafficking machinery, two factors drive subcellular protein localization within the micron-scale confines of the bacterial cytoplasm. First, proteins that are synthesized in the cytosol move simply by diffusion (either in three dimensions in the cytosol for soluble or peripheral membrane proteins, or in two dimensions in the membrane for integral membrane proteins) (13). Second, proteins are captured at the correct subcellular location by a chemical or physical cue that uniquely defines that location in the cell (14). Most often, the localization cue is a pre-localized protein, but physical cues such as the local curvature of the membrane or proximity to another cellular compartment can also serve as subcellular localization cues (15–23). Identifying the terminal cues that ultimately drive subcellular protein localization in bacteria, especially those for the localization of integral membrane proteins, is a longstanding challenge in bacterial cell biology.

Subcellular protein localization has been extensively studied in *Bacillus subtilis*, especially during the process of spore formation (sporulation), which involves extensive membrane reorganization and the creation of different subcellular compartments that provide the opportunity to study different types of protein localization events in a single model system (24, 25) (Fig. 1A). Sporulation initiates in the rod-shaped *B. subtilis* by asymmetric cell division, which generates two compartments that initially lie side-to-side: a smaller forespore that will eventually mature into a dormant spore and a larger mother cell that nurtures the developing forespore and will eventually lyse. Next, the mother cell engulfs the forespore, generating a cell-within-a-cell configuration in which the forespore is enveloped by a double membrane envelope and is a spherical, organelle-like compartment inside the mother cell cytosol (26). The mother cell then constructs a peptidoglycan “cortex” between the double membrane envelope and a proteinaceous “coat” atop the outer forespore membrane (27, 28). Coat assembly initiates with the localization of a peripheral membrane protein that preferentially embeds in the convex outer forespore membrane and recruits a structural protein that forms the coat basement layer (29). Although assembly of the spore coat represents the most significant protein localization event around the forespore with ∼80 proteins localizing in an ordered fashion, several integral membrane proteins that are required for cortex assembly also localize to the mother cell face of the asymmetric septum (later the outer forespore membrane), whose mechanisms of localization are poorly understood.

**Figure 1.**
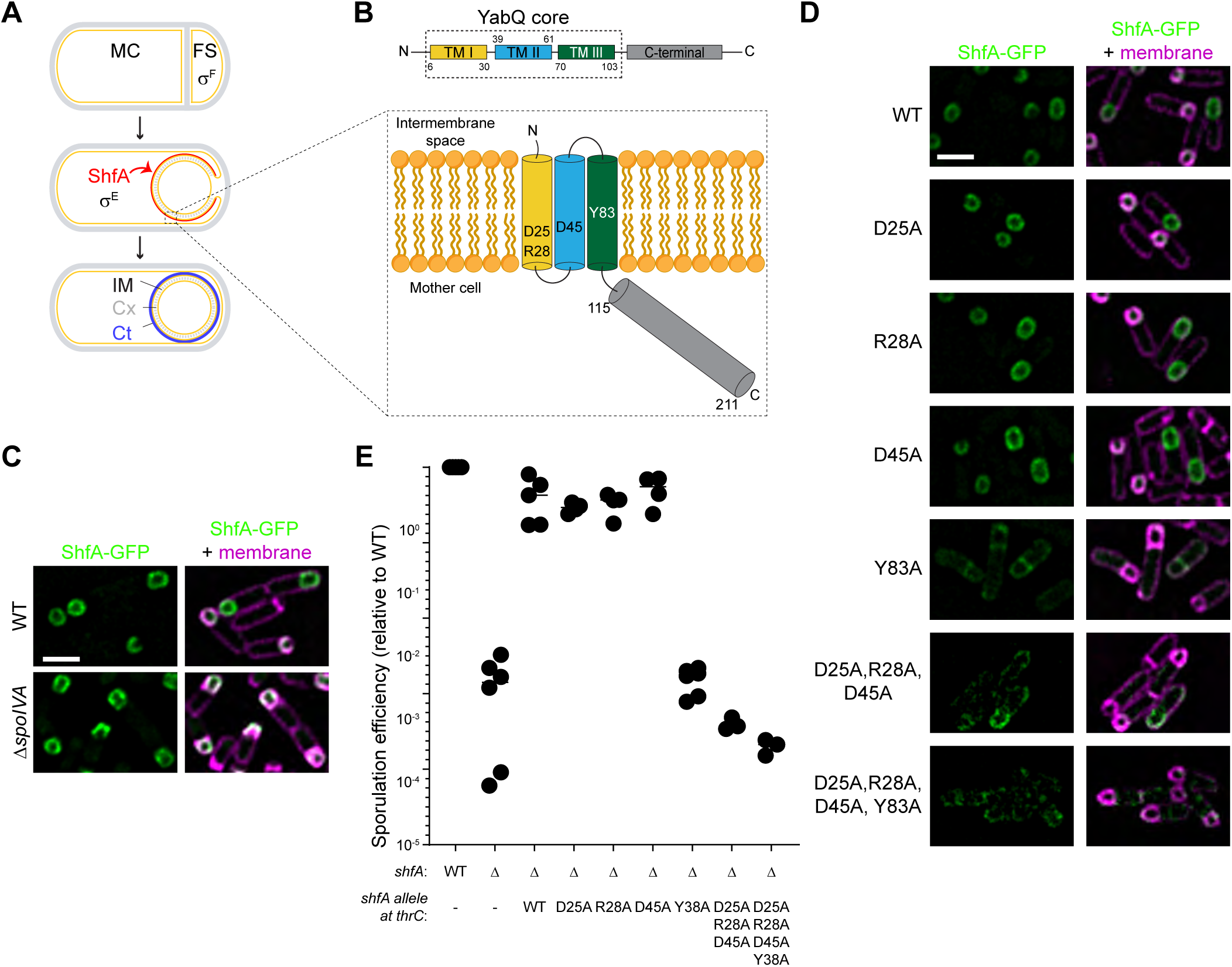
Polar residues in the ShfA transmembrane domain direct subcellular localization of ShfA. (A) Schematic of sporulation in *B. subtilis*. Cell wall depicted in gray; membranes, yellow. Nutrient depletion drives asymmetric division of the progenitor cell, resulting in two daughter cells: a large mother cell (MC) and a smaller forespore (FS). The transcription factor σ^F^ is specifically activated in the forespore after which σ^E^ is specifically activated in the mother cell. As the mother cell engulfs the forespore, ShfA is produced under control of σ^E^ and localizes to the outer forespore membrane. After engulfment, the spore coat (Ct, blue) assembles on the outer forespore membrane. The spore cortex (Cx) cell wall is then constructed in the intermembrane space (IM) between the two membranes encasing the forespore. (B) Schematic of the ShfA domain architecture and predicted membrane topology. The YabQ core consists of three transmembrane segments embedded in the outer forespore membrane in an N-out orientation with the C-terminal domain in the mother cell cytosol. (C) Subcellular localization of ShfA-GFP 3.5 h after induction of sporulation in WT cells and in the absence of the spore coat basement layer protein SpoIVA. Left: fluorescence from ShfA-GFP (green); right: overlay, GFP and membranes visualized with FM4-64 (pink). Strains: VP375, VP376. Scale bar: 2 µm. (D) Subcellular localization of WT ShfA-GFP or variants harboring alanine substitutions of indicated polar residues in the YabQ domain. left: fluorescence from GFP (green); right: overlay, GFP and membranes (pink). Strains: CW205, CC10, CC11, CC12, CC13, CC30 and VP069. Scale bar: 2 µm. (E) Sporulation efficiencies relative to WT, determined by resistance to wet heat, of strains harboring the indicated allele of *shfA*. *thrC* is a chromosomal locus in which different alleles of *shfA* are inserted. Bars represent mean; data points indicate independent experiments.

One such integral membrane protein is ShfA (YabQ), a broadly conserved sporulation protein that is produced in the mother cell and localizes to the outer forespore membrane, where it is required for cortex assembly (Fig. 1A) (30, 31). ShfA is part of the ShfA-ShfP (YvnB) pathway wherein *B. subtilis* cells in a population that initiate sporulation early signal to other cells in the population (via extracellular glycerol) to delay that population’s entry into sporulation (32). The secreted glycerol is somehow toxic to the sporulating cells that secrete it, and ShfA acts as an “antidote” to the apparent sporulation inhibitory activity of the secreted glycerol.

The antidote activity of ShfA requires its proper localization to the forespore surface. Here, we investigate the mechanism by which ShfA recognizes this specific patch of membrane in the mother cell cytosol. We show that ShfA has an unusual membrane topology and that it localizes to the correct membrane, not by diffusing laterally in the membrane upon aided insertion (like previously described bacterial integral membrane proteins), but by spontaneously inserting into the preferred membrane. The localization cue is not mediated by local membrane curvature, but instead a broadly conserved feature of division septa that requires active peptidoglycan synthesis. We demonstrate that the chemical cue is undecaprenyl phosphate (UndP), the bacterial isoprenoid lipid carrier that traffics polysaccharides across membranes (33), and that UndP likely directly drives the targeted insertion of ShfA into the outer forespore membrane.

## RESULTS

### The YabQ transmembrane domain is sufficient for ShfA function and subcellular localization

ShfA is produced in the mother cell compartment under control of the sporulation-specific σ^E^ sigma factor and localizes to the outer forespore membrane (Fig. 1A), and deletion of *shfA* results in a severe sporulation defect (30, 31). Interestingly, while ShfA homologs are found only in sporulating Bacillota (formerly Firmicutes), they exhibit great variability in length (Fig. S1A). Hence, to define its conserved functional core, we comprehensively collected ShfA homologs and constructed a multiple sequence alignment, which we used to dissect its three-dimensional structure predicted with AlphaFold3 and its transmembrane topology predicted with Phobius and DeepTMHMM. These predictions revealed that the conserved core of the ShfA family was a unique N-terminal domain comprised of 3 transmembrane helices with an unusual packing followed by a C-terminal cytoplasmic coiled-coil tail (Figure 1B). We named the conserved core of the ShfA family the YabQ domain, after the original name of the family. In many species of Bacillota, including *B. subtilis*, this cytoplasmic coiled-coil segment of the YabQ domain leads into a highly variable helical amphipathic extension, the variation in whose length accounts for the observed length variability in ShfA homologs (Fig. S1A).

Both the Phobius and DeepTMHMM predictions indicated that the YabQ domain is embedded in the membrane with an extracytoplasmic N-terminus followed by the three transmembrane helices and the C-terminal region in the cytoplasm. While this is an unusual orientation, it nonetheless satisfies the “positive inside rule” which states that positively charged residues are more abundant in cytoplasmic regions of transmembrane proteins (34–36) (Fig. 1B). To test this predicted membrane topology, we generated a series of C-terminal truncations fused to a *phoA-lacZα* dual reporter, wherein PhoA is only active when extracytosolic and β-galactosidase is only active in the cytoplasm (37) and expressed the constructs in *E. coli*. The activities of each reporter were consistent with the YabQ domain of ShfA lacking a signal sequence (36) and ShfA being oriented in the membrane with an extracytosolic N-terminus and a cytosolic C-terminus (Fig. S1B).

The sporulation-specific protein SpoIVA is an ATPase that is anchored to the outer forespore membrane and directly or indirectly recruits ∼80 different proteins that form the spore coat (6, 29, 38). To determine whether ShfA is a spore coat protein, we examined its localization in the absence of SpoIVA. Deletion of *spoIVA* did not abrogate localization of ShfA-GFP to the forespore surface (Fig. 1C), indicating that ShfA localizes to the forespore in a SpoIVA-independent manner.

The YabQ domain is marked by a set of seven well-conserved polar residues. While three of these are predicted to lie in the extracytoplasmic loops, four of them (D25, R28, D45, and Y83) are located in the transmembrane helices (Fig. 1B). These residues are predicted to cluster in a putative ligand-binding pocket near the cytoplasmic leaflet of the outer forespore membrane. Substituting any of the three charged residues with Ala did not affect the localization of ShfA-GFP during sporulation, but substituting Y83 alone, substituting all three charged residues simultaneously, or substituting all four polar residues with Ala resulted in the promiscuous localization of ShfA-GFP to the mother cell membrane (Fig. 1D). Consistent with the mis-localization patterns, complementation of the Δ*shfA* strain with variants harboring single substitutions of each charged residue did not result in a sporulation defect in vivo (Fig. 1E, lanes 1-6). However, strains expressing *shfA*^Y83A^, *shfA*^D25A,R28A,D45A^, or *shfA*^D25A,R28A,D45A,Y83A^ exhibited a ∼1000-fold reduction in sporulation efficiency (Fig. 1E, lanes 7-9), even though the variants were produced at similar levels as WT ShfA (Fig. S1E). Since ShfA orthologs exhibit limited conservation and high length variability in the cytosolic C-terminal tail (Fig. S1A), we predicted that the cytosolic tail is unlikely to play a role in the function of ShfA (Fig. S1A). We assessed this prediction by generating a C-terminal truncation of ShfA to produce ShfA^1-112^-GFP, which retains just the core YabQ domain as defined by our sequence and structure analysis. The truncation did not affect localization of the protein to the forespore surface (Fig. S1C). Notably, complementing the Δ*shfA* strain with just *shfA*^1-114^ or *shfA*^1-103^ fully restored sporulation efficiency (Fig. S1D, lanes 1-5). Further truncation of the YabQ domain to retain just the 3 transmembrane helices (ShfA^1-86^-GFP) supported forespore localization but did not complement the sporulation defect of Δ*shfA*, and any further truncation of the YabQ domain resulted in mis-localization of the protein and abrogated function (Fig. S1C-D), likely due to reduced stability (Fig. S1E). Thus, the YabQ domain is necessary and sufficient for ShfA localization and function. These results also suggest that the correct subcellular localization of ShfA depends specifically on the three transmembrane helices of the YabQ domain, which contains four conserved intramembrane polar residues.

### ShfA undergoes directed insertion into the outer forespore membrane

Bacterial integral membrane proteins typically achieve their proper subcellular localization by a “diffusion-and-capture” mechanism, wherein the protein is first (actively) inserted into the membrane, followed by lateral diffusion in the membrane until it is captured by a subcellular landmark (typically a pre-localized protein) (14, 39). In contrast, peripheral membrane proteins, which exhibit an on- and off-rate from membranes and therefore sample multiple locations, can undergo “directed insertion” into a preferred membrane. This mechanism, for example, is exploited by proteins that preferentially embed into membranes with specific curvature (18, 19, 29, 40). To distinguish between these models for ShfA localization, we examined ShfA localization at a late time point during sporulation after engulfment completes and the forespore topologically detaches from the mother cell. The completion of engulfment may be monitored by using two different fluorescent membrane dyes: a membrane-permeable dye that stains all membranes, and a membrane-impermeable dye that cannot stain the forespore after the completion of engulfment (14, 41). As previously reported, after the completion of engulfment (evidenced by the inability of the dye-accessible membrane to detect the forespore; Fig. 2A) the integral membrane protein SpoIVFB localized to membrane surrounding the mother cell but was largely unable to localize to the forespore surface (Fig. 2A (14)). For unknown reasons, the outer forespore membrane is unable to support the insertion of an integral membrane protein after the completion of engulfment, presumably because the protein translocation machinery is either absent or not functional at that location (14). In contrast, nearly half of the cells exhibited proper ShfA-GFP localization even after the completion of engulfment (Fig. 2A), suggesting that ShfA is able to localize to the forespore via a mechanism independent of diffusion-and-capture.

**Figure 2.**
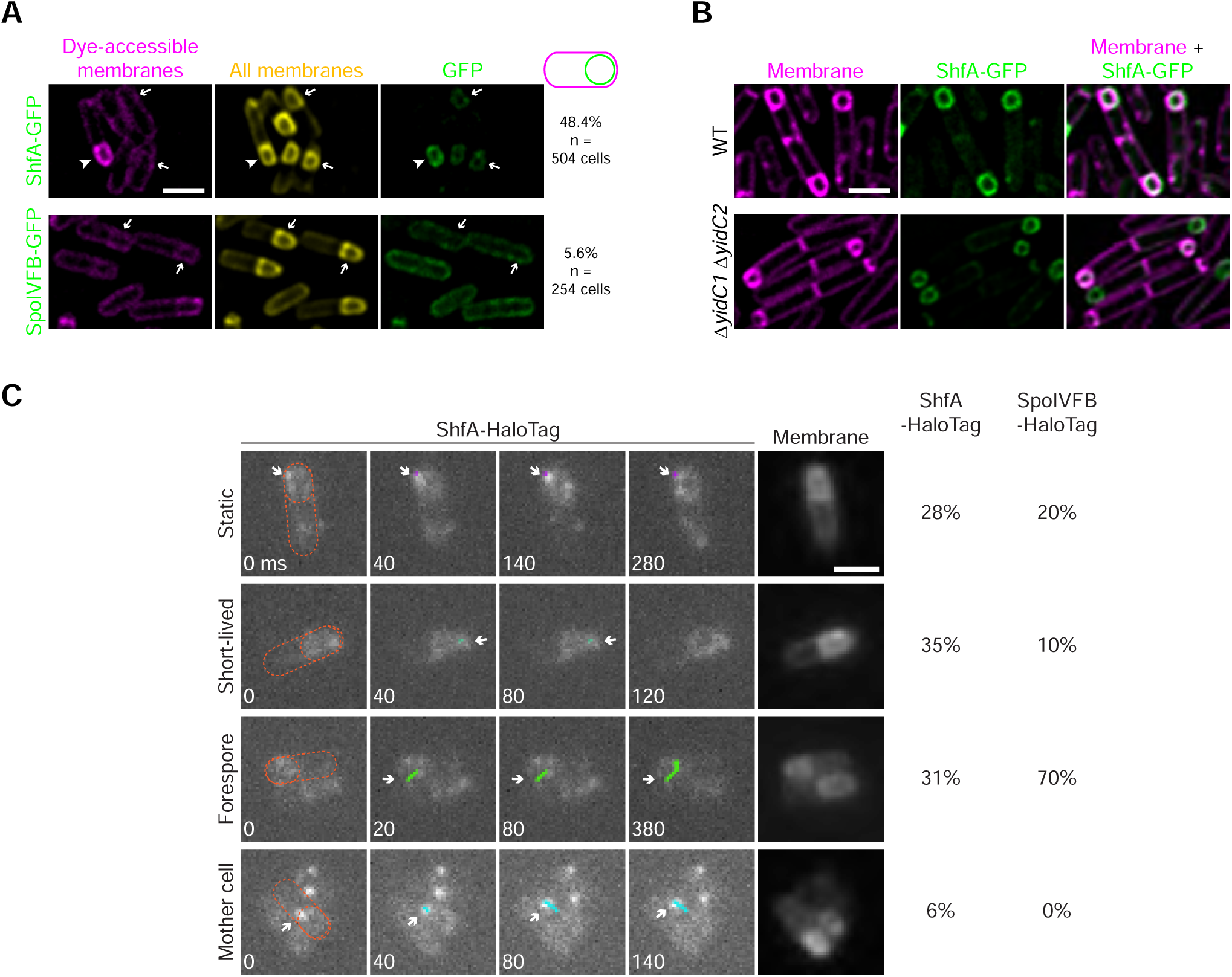
Directed insertion of ShfA onto the outer forespore membrane in a *yidC*- and σ^F^-independent manner. (A) Localization of ShfA-GFP (top) or SpoIVFB-GFP (bottom) produced after topological isolation of the forespore. The membrane surrounding fully engulfed forespores was stained by a membrane-permeating dye (Mitotracker Deep Red; middle, yellow) but not by a nonpermeating dye (FM4-64; left, pink). Right: fluorescence from GFP (green). Arrows indicate fully engulfed forespores; arrowhead indicates a forespore that is not engulfed. Strains: CW189 and BDR647 (B) Localization of ShfA-GFP in the presence (top) or absence (bottom) of the membrane insertases YidC1 and YidC2 3.5 h after induction of sporulation. Strains: CW205 and VP298. (C) Columns 1-4: representative images of single molecule trajectory tracking of ShfA-HaloTag on forespores using JFX554 dye. Time (ms) of each frame relative to the first frame is indicated in each frame. Outline of an individual cell of interest, determined by the cells visualized with membrane stain MitoTracker Deep Red FM (column 5), is indicated using an orange dashed line in the first frame. Individual trajectories are classified as “mobile, forespore” (mobile within the forespore boundary; green), “mother cell” (mobile trace initiating within the forespore but moving outside of the forespore boundary into the mother cell; cyan), or “immobile forespore” (relatively short track length within the forespore; pink). The percentage of observed trajectories for ShfA-HaloTag and SpoIVFB-HaloTag (n = 154 and n = 132 trajectories, respectively, from >30 cells) displaying each behavior is indicated to the right. Strains: VP461 and VP462. Scale bars: 2 µm.

Integral membrane proteins that lack a canonical signal sequence and exhibit an N-out orientation in the membrane, including those that historically were proposed to insert spontaneously, usually require the YidC insertase for membrane insertion, which can work independently or in concert with the SecEYG channel (42–44). *B. subtilis* encodes two YidC orthologs, the sporulation-specific YidC1 (SpoIIIJ) and YidC2 (YqjG), both of which have been implicated in membrane protein biogenesis and secretion (45–47). Deleting both *yidC1* and *yidC2* did not block entry into sporulation and did not impair localization of ShfA-GFP to the forespore surface (Fig. 2B). Together with the previous observation that the forespore surface does not support Sec-mediated insertion of integral membrane proteins after the completion of engulfment (14), these results suggest that ShfA spontaneously inserts into the outer forespore membrane without requiring a dedicated insertase.

A model that invokes the unaided membrane insertion of ShfA would predict that ShfA, unlike a typical integral membrane protein that remains tethered on a membrane, likely samples multiple sites within a cell, similar to a peripheral membrane protein (29). To test this prediction, we performed in vivo single-molecule imaging of ShfA fused to HaloTag and, as a control for an embedded transmembrane protein, also examined the intracellular motion of SpoIVFB-HaloTag (48) in conjunction with a membrane-permeable dye that covalently binds to the HaloTag fusion (49). In the presence of excess dye, ShfA-HaloTag and SpoIVFB-HaloTag localized preferentially to the forespore membrane, indicating that the tag did not interfere with the proper localization of either protein (Fig. S2A). In the presence of limiting amounts of dye, we observed the appearance and disappearance of single molecules of both fusion proteins. Based on single molecule trajectories computed for both proteins on the forespore, we classified them into three categories: “mobile, forespore”, in which a single molecule moved exclusively within the boundary of the forespore; “mother cell” in which a single molecule was visualized as starting within the forespore boundary and then moving outside of it into the mother cell; and “immobile, forespore”, in which the single molecule signal displayed a relatively short track length in the forespore (Fig. 2C). The entirety of observed SpoIVFB-HaloTag single molecule trajectories (n = 132 trajectories from >30 different cells) were classified as either “mobile, forespore” or “immobile, forespore” and we observed no trajectories in which SpoIVFB-HaloTag left the forespore and entered the mother cell (Fig. 2C). In contrast, we observed that 6% of ShfA-HaloTag trajectories moved from the forespore to the mother cell cytosol (n = 154 trajectories from >30 different cells). Combined with the observation that ShfA, but not SpoIVFB, can localize to the forespore after engulfment (Fig. 2A), the data in sum suggest the ability of ShfA to bind to the forespore membrane and subsequently dissociate from it.

### ShfA localization is mediated by a widely conserved septal feature

The proper localization of at least one integral membrane protein during sporulation in *B. subtilis* (SpoIIIAH), which is produced in the mother cell and localizes on the mother cell face of the polar septum before engulfment initiates, requires direct interaction with the extracellular domain of a protein produced in the forespore compartment (21, 22). To test if ShfA can localize to a flat polar septum, we first examined ShfA-GFP localization in a previously characterized mutant cell that is arrested during sporulation before the onset of engulfment (50). In this mutant background, ShfA-GFP localized exclusively to the flat polar septum (Fig. S2B). Additionally, similar to the requirement for proper ShfA localization to the outer forespore membrane, localization of ShfA-GFP to the flat septum was mediated by the YabQ domain of ShfA and the four conserved intramembrane polar residues in the YabQ domain (Fig. S2B). To test if forespore-produced proteins are required for proper localization, we deleted the gene encoding the first forespore-specific sigma factor (σ^F^) and genetically bypassed σ^F^ requirement for progression through sporulation by producing σ^E^ directly under control of the master transcriptional regulator for entry into sporulation (Spo0A). In the absence of σ^F^, ∼41% of cells that displayed a GFP signal still formed a polar septum to which ShfA-GFP localized (Fig. 3A, top row). Interestingly, approximately 34% of these cells instead harbored a septum at mid-cell to which ShfA-GFP also localized. The results therefore indicate that ShfA preferentially inserts into the mother cell face of the polar septum independently of any factors produced in the forespore but also raised the possibility that ShfA may indiscriminately localize to any cell division septum.

**Figure 3.**
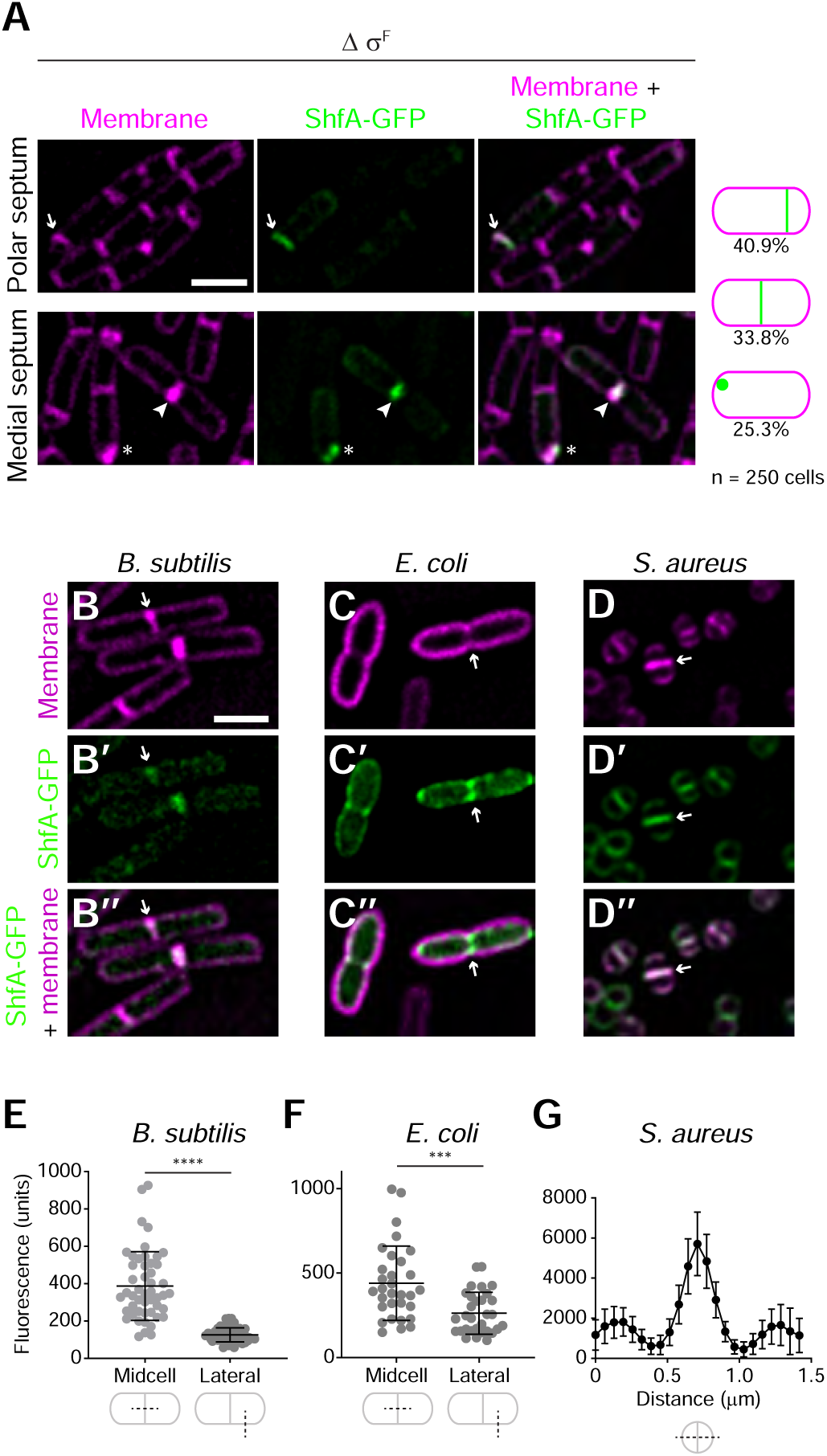
ShfA subcellular localization cue is broadly conserved. (A) Localization of ShfA-GFP in a strain that bypasses σ^F^ activation. Top row: localization at polar septum; bottom row: localization at an aberrant medial septum during sporulation. Strain VP010. Left: membranes (pink); middle: fluorescence from ShfA-GFP (green); right: overlay, membranes and GFP. (B-D’’) Localization of ShfA-GFP produced upon induction during exponential growth in (B-B’’) *B. subtilis* (strain VP050), (C-C’’) *E. coli* (strain BL21(DE3) pVP008), and (D-D’’) *S. aureus* (strain RN4220 pVP012). (B-D) Membranes (pink); (B’-D’) fluorescence from ShfA-GFP (green); (B’’-D’’) overlay, membrane and GFP. Arrows indicate division septa. Scale bars: 2 µm. (E-F) Quantification of GFP fluorescence across division septa (mid-cell) or lateral membrane (lateral) in (E) *B. subtilis* (n > 250 cells) and (F) *E. coli* (n >150 cells), or (G) across the width of a dividing *S. aureus* cell (n > 250 cells). In (E-F) bars indicate mean; errors: S.D. **** indicates *p* < 0.0001; *** indicates *p* < 0.0002 (t-test). In (G) data points represent mean; errors: S.D.

We therefore tested if a normal division septum, even in the absence of sporulation-specific factors, could recruit ShfA. Indeed, ShfA-GFP artificially produced during vegetative growth in *B. subtilis* localized preferentially to the division septum (Fig. 3B, E). We next tested if ShfA would localize to the division septum of the Gram-negative bacterium *E. coli*. During exponential growth, ShfA-GFP fluorescence was enriched at the division septum, with additional diffuse localization along the cell periphery (Fig. 3C, F). Finally, we examined ShfA-GFP localization in the Gram-positive *Staphylococcus aureus*, a non-sporulating member of the Bacillota lineage which, although more closely phylogenetically related to *B. subtilis* than *E. coli*, is spherical and harbors a pentaglycine crossbridge that links adjacent peptide stems in its peptidoglycan (unlike the rod-shaped *B. subtilis* and *E. coli* in which meso-diaminopimelic acid directly crosslinks with D-alanine in an adjacent peptide stem (51)). Similar to its localization pattern in *B. subtilis* and *E. coli*, ShfA-GFP preferentially localized to division septa when produced in *S. aureus* (Fig. 3D, G), indicating that ShfA-GFP recognizes a feature of the division septum that is broadly conserved in bacteria.

### Inhibition of peptidoglycan synthesis abrogates ShfA localization

Elaborating the division septum involves active cell wall remodeling that drives membrane constriction (52, 53). To test if ShfA localization to vegetative division septa in *B. subtilis* requires active cell wall synthesis, we inhibited cell wall synthesis using either vancomycin, which blocks peptidoglycan crosslinking (54) or tunicamycin, which inhibits the initial steps of wall teichoic acid synthesis (55). Inducing expression of *shfA-gfp* during vegetative growth resulted in preferential localization of ShfA-GFP at the division septum (Fig. 4A’’) at sites coincident with active peptidoglycan synthesis (as determined by incorporation of the fluorescent D-amino acid 7-hydroxycoumarincarbonylamino-D-alanine (HADA); Fig. 4A’) and membrane constriction, evidenced by increased membrane staining (Fig. 4A’’). When vancomycin was added at the time of ShfA-GFP induction and imaged 30 min later, septum-localized ShfA-GFP was absent in 72% of cells (Fig. 4B-B’’). In contrast, ShfA-GFP continued to localize robustly in cells that were similarly treated with tunicamycin (Fig. 4C-C’’), suggesting that a factor related to synthesis of the peptidoglycan component of the cell wall, but not the teichoic acid component, mediates ShfA localization to division septa. Vancomycin targets the terminal D-Ala-D-Ala residues of the pentapeptide stem, thereby preventing transpeptidation that results in reduced peptidoglycan cross-linking. To examine if peptidoglycan cross-linking mediates ShfA localization, we examined ShfA-GFP localization in a strain lacking the carboxypeptidase DacA, which fails to cleave the terminal D-Ala residue of the pentapeptide stem, resulting in increased vancomycin resistance (56, 57). Indeed, a recent study revealed that the *Streptococcus pneumoniae* cell division protein MapZ preferentially binds to DacA-modified peptidoglycan (58). However, unlike the behavior of MapZ, in a Δ*dacA* strain ShfA-GFP continued to localize to division septa that also displayed active incorporation of the HADA dye and membrane invagination (Fig. 4D-D’’). Furthermore, deleting sporulation-specific peptidoglycan synthesis genes (*dacB*, *spoVB*, *yabM*, *spoVE*, and *spoVD*) also did not alter ShfA localization during sporulation (Fig. S3). We therefore conclude that while ShfA-GFP localizes to sites of active peptidoglycan synthesis, its localization is not dependent on an interaction with known peptidoglycan-biogenesis proteins or a chemical cue that is present on mature peptidoglycan.

**Figure 4.**
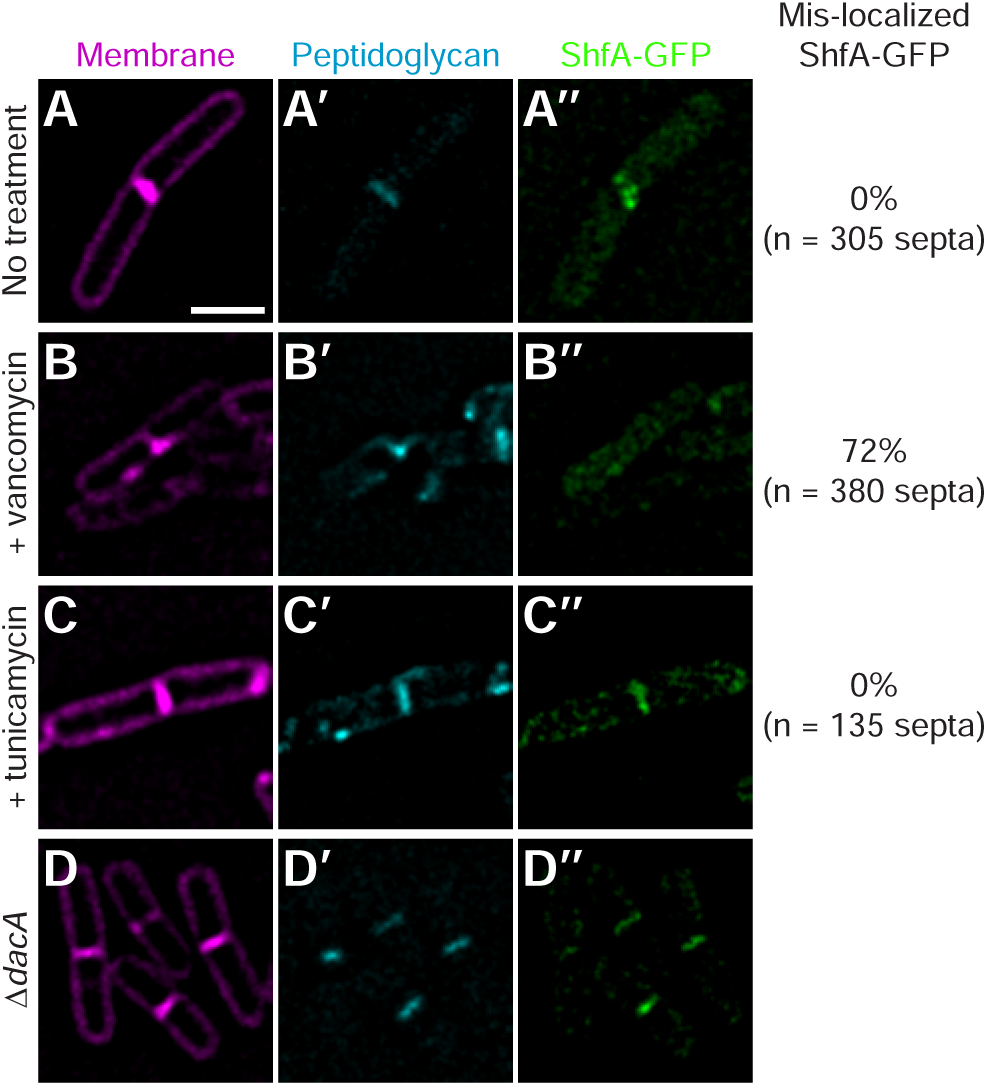
Disruption of active peptidoglycan synthesis alters ShfA localization. Subcellular localization of ShfA-GFP in vegetatively growing *B. subtilis* after (A-A’’) no treatment, (B-B’’) 0.5 µg ml^-1^ vancomycin treatment, or (C-C’’) 16 µg ml^-1^ tunicamycin treatment, or (D) in the absence of *dacA*. (A-D) Membranes visualized with FM4-64; (A’-D’) fluorescence from HADA incorporation into peptidoglycan; (A’’-D’’) fluorescence from ShfA-GFP. Frequency of ShfA-GFP mis-localization under each condition is reported to the right. Scale bar: 2 µm.

### Active peptidoglycan synthesis drives ShfA localization

DivIC (FtsB in *E. coli*) is a component of the cell division machinery and forms a regulatory subcomplex with two other proteins that participate in the recruitment and stabilization of peptidoglycan synthases at the division septum during cell division (59–61). Interestingly, the *divIC* gene is part of a conserved gene neighborhood seen across at least 89% of the sporulating Bacillota, where it lies immediately downstream of *shfA*, as in *B. subtilis*. The *shfA*-*divIC* gene neighborhood association spans all major sporulating lineages, including *Bacillus*, *Paenibacillus*, *Halobacillus*, *Clostridium*, as well as more divergent genera such as *Sporosarcina*, *Heliobacterium* and *Natranaerobius*, suggesting a strong functional coupling between these proteins. We therefore examined whether DivIC activity at the division septum during cell wall synthesis drives ShfA localization. In actively dividing *B. subtilis*, 69% of nascent septa displayed sfGFP-DivIC localization, whereas only 31% of apparently mature septa displayed sfGFP-DivIC localization (Fig. 5A; red arrows and arrowheads, respectively). We next tested whether ShfA localization depends on DivIC activity using a previously described temperature-sensitive *divIC* allele (62). In otherwise WT cells, ShfA-GFP localized at division septa, at sites which also displayed evidence of both active peptidoglycan assembly (evidenced by HADA incorporation) and membrane invagination (evidenced by increased FM4-64 staining) (Fig. 4B). In contrast, cells harboring the temperature-sensitive *divIC* allele grown at the nonpermissive temperature displayed 1) fewer division septa, which resulted in cell elongation; and 2) aberrant division septa that showed either sites of increased peptidoglycan incorporation but no concomitant membrane invagination, or sites where the membrane invaginated with no concomitant peptidoglycan incorporation. We exploited this uncoupling of membrane invagination and peptidoglycan synthesis in the *divIC*^ts^ mutant to examine ShfA localization to each of these sites. As expected, ∼99% of sites where we detected both membrane invagination and peptidoglycan incorporation also showed ShfA-GFP colocalization (Fig. 5B, white arrows). However, when only membrane invagination occurred, ShfA-GFP colocalization was reduced to ∼33%, and no ShfA-GFP localization was detected at sites where only peptidoglycan incorporation occurred (Fig. 5B, gray arrows), indicating that ShfA-GFP localization requires the concerted events of both peptidoglycan assembly and membrane invagination.

**Figure 5.**
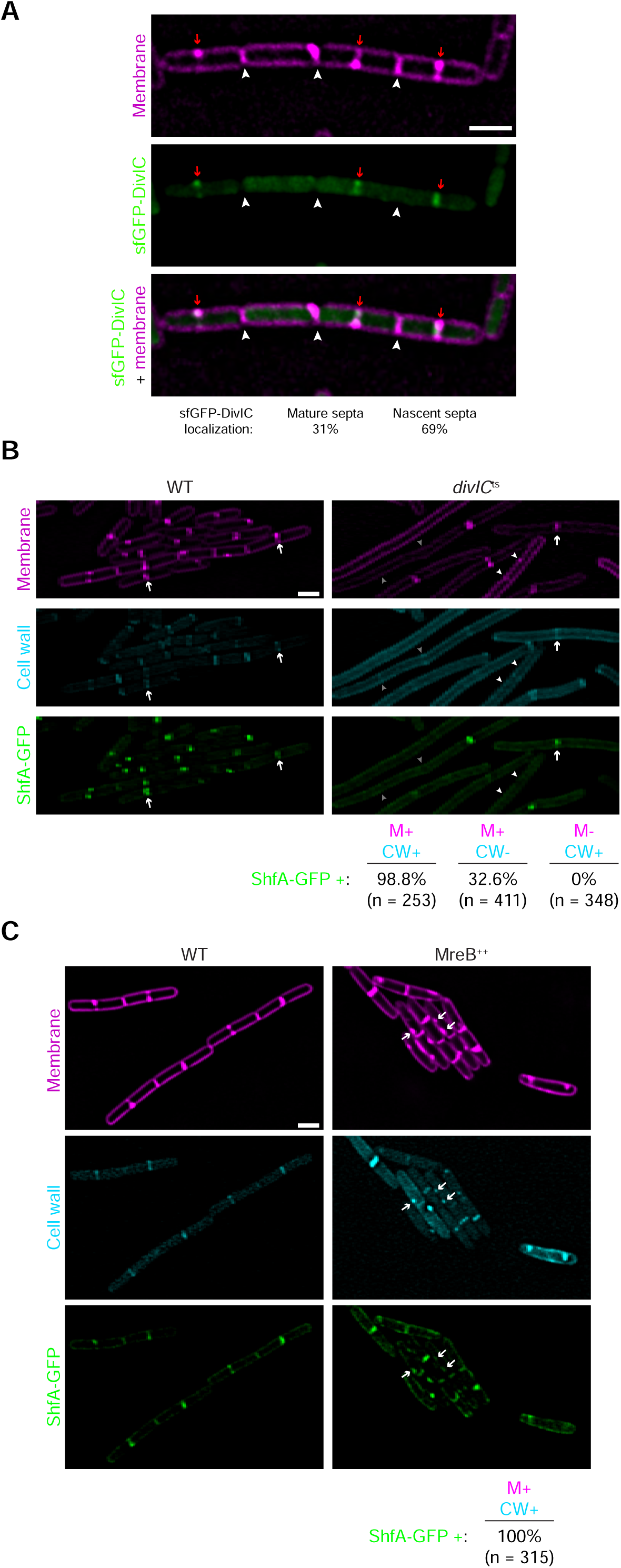
ShfA co-localizes preferentially to sites of active peptidoglycan synthesis, not simply membrane invagination. (A) Localization of sfGFP-DivIC in vegetatively growing *B. subtilis*. Top: membranes visualized with FM4-64; middle: fluorescence from sfGFP-DivIC; bottom: overlay, membrane and sfGFP. Red arrows indicate nascent division septa; white arrowheads indicate mature septa. Localization frequency of sfGFP-DivIC at mature septa or nascent septa indicated below. (B) Localization of ShfA-GFP in the (left) presence or (right) absence of *divIC*. Top: membranes visualized with FM4-64; center: fluorescence from HADA incorporation into peptidoglycan; bottom: fluorescence from ShfA-GFP. Arrows indicate sites of membrane invagination and HADA incorporation; white arrowheads indicate sites of membrane invagination but no HADA incorporation (and no fluorescence from GFP); gray arrowheads indicate sites of HADA incorporation but no membrane invagination. (C) Localization of ShfA-GFP in (left) otherwise WT cells or (right) cells overproducing MreB during vegetative growth. Top: membranes visualized with FM4-64; center: fluorescence from HADA incorporation into peptidoglycan; bottom: fluorescence from ShfA-GFP. Arrows indicate sites of aberrant HADA incorporation and membrane invagination that coincide with GFP fluorescence. Strains: CW189, VP350, VP433. Scale bars: 2 µm.

We then investigated if increasing peptidoglycan synthesis along the lateral edge of the cell could redirect ShfA localization away from the division septum and towards those aberrant sites. To this end, we overproduced MreB during vegetative growth, which has previously been shown to alter cell shape as a result of spurious peptidoglycan assembly at incorrect sites (63). Cells harboring an additional copy of *mreB* at an ectopic locus displayed an abnormal morphology, including aberrant sites of membrane invagination along the lateral edge that coincided with high levels of peptidoglycan assembly (Fig. 5C). At 100% of these sites that we observed (n = 315) we also observed the colocalization of ShfA-GFP (Fig. 5C), suggesting that ShfA targeting is governed primarily by zones of active peptidoglycan assembly that causes a perturbation in the membrane, and not necessarily by a unique feature of the division septum.

### ShfA requires undecaprenyl phosphate for proper localization

The structure of the YabQ domain predicted using AlphaFold3 revealed an unusual packing of the three TM helices, with helices 1 and 2 lying parallel to each other and helix 3 lying across the surface of the former two (plDDT> 90). As a result, these three helices form a distinct intramembrane pocket, whose floor is partly closed by the cytoplasmic coiled-coil extension (Fig. 6A). The broader end of this pocket, directed toward the cytoplasmic leaflet of the lipid bilayer, houses the four conserved polar residues, which are critical for ShfA localization and function (Fig. 1D). Hence, the pocket exhibits an amphipathic character, with a cytoplasm-facing polar region and a membrane-spanning hydrophobic region. Accordingly, we reasoned that the YabQ domain might specifically bind a lipid, with its head group localized near the pocket’s polar region and its aliphatic tail shielded by the pocket’s hydrophobic walls. Given our observation that ShfA localizes to membrane sites associated with active peptidoglycan synthesis, we wondered if ShfA localization may directly depend on undecaprenyl phosphate (C55-P), the carrier molecule for peptidoglycan across the cytoplasmic membrane, that forms the transmembrane portion of the lipid I and lipid II peptidoglycan precursors (64). We accordingly tested this *in silico* by creating an AlphaFold3 model of the YabQ domain with undecaprenyl diphosphate. The lipid was indeed bound in the pocket of the YabQ domain with the conserved polar residues interacting with the phosphates in the head group (Fig. 6B). This suggested that direct intramembrane binding of undecaprenyl lipids by the YabQ domain might be key to the localization and function of ShfA.

**Figure 6.**
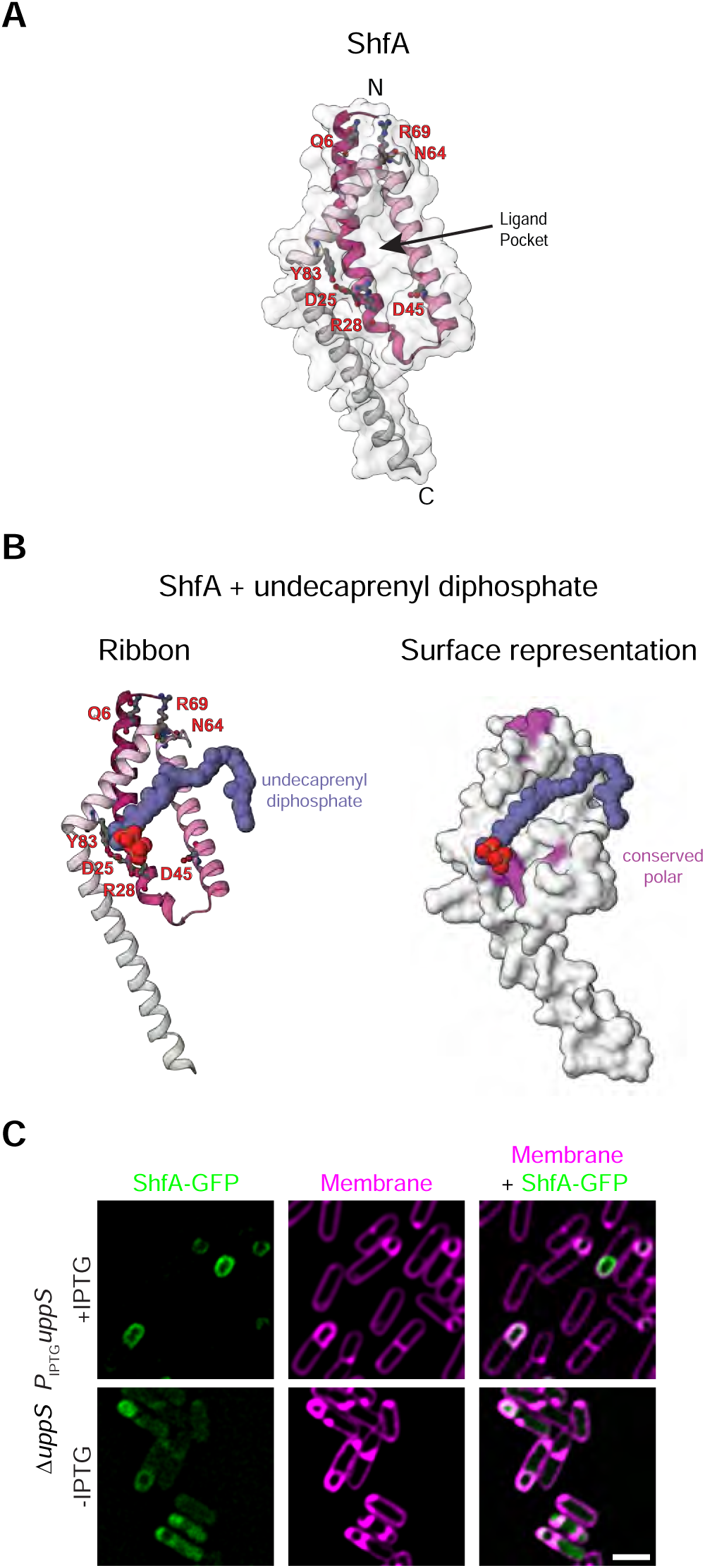
Depletion of undecaprenyl phosphate disrupts ShfA localization to the forespore surface. (A) AlphaFold3-predicted structure of the conserved YabQ core of ShfA. The model includes a translucent surface view; predicted residues in the ligand-binding pocket are indicated and shown as ball-and-stick models. (B) AlphaFold3-predicted structure of the YabQ domain of ShfA bound to undecaprenyl diphosphate. Left: ribbon diagram showing predicted ligand-binding residues represented as ball-and-stick models; undecaprenyl diphosphate depicted in purple. Right: surface representation with conserved polar residues highlighted in magenta. (C) Subcellular localization of ShfA-GFP in the (top) continued presence and (bottom) depletion of IPTG, which induces *uppS* (encoding undecaprenyl pyrophosphate synthetase) 2.5 h after induction of sporulation. Left: fluorescence from ShfA-GFP (green); center: membranes visualized with FM4-64 (pink); right: overlay, GFP and membranes. Strain VP422. Scale bar: 2 µm.

We therefore constructed a strain in which *uppS*, which encodes undecaprenyl pyrophosphate synthetase that provides the precursor for UndP synthesis, was expressed during sporulation under the control of an inducible promoter in a cell harboring a deletion of *uppS*. In the presence of inducer, cells continued through sporulation, and ShfA-GFP localized to the forespore surface (Fig. 6C). However, removing the inducer during sporulation resulted in the mis-localization of ShfA-GFP, indicating that proper localization of ShfA indeed requires UndP. Taken together, the data are consistent with a model in which UndP is a chemical cue that recruits ShfA to sites of active peptidoglycan synthesis.

## DISCUSSION

In eukaryotes, precise spatial and temporal localization of integral membrane proteins is carried out by complex sorting and trafficking systems that direct specific proteins across tens of microns to chemically distinct organelles (65–68). In bacteria, proteins are sorted to specific subcellular regions, even in the typical absence of organelles, to achieve a highly organized cytoplasm at the nanometer or micron scale. In the present study, we examined the subcellular localization of a sporulation-specific integral membrane protein, ShfA, in *B. subtilis*, which localizes to the asymmetric division septum and later remains associated with the outer forespore membrane after engulfment. We report that the N-terminal transmembrane YabQ domain of ShfA, which displays an unusual N-out membrane topology (35) and harbors four conserved intramembrane polar residues, is necessary and sufficient for the proper localization of ShfA. While most transmembrane proteins are actively inserted into the membrane by the SecEYG machinery and the insertase YidC, we report that YabQ domain of ShfA likely directs its spontaneous insertion into the membrane. Curiously, ShfA membrane insertion occurred preferentially at sites of active peptidoglycan synthesis, such as division septa. In silico modeling suggests that the YabQ domain of ShfA harbors a binding pocket whose hydrophobic walls can accommodate the C55 undecaprenyl moiety, while a cluster of four polar amino acids at its base can accommodate the phosphate group. Consistent with this, our genetic evidence suggests that the universal bacterial lipid carrier UndP, which traffics polysaccharides across membranes and is required for peptidoglycan synthesis, is the subcellular localization cue for ShfA.

This localization mechanism appears distinct from other strategies used by membrane proteins to localize to division septa in bacteria. For example, proteins containing LysM, the bacterial SH3, or SPOR domains bind distinct features of peptidoglycan, including denuded glycan strands found at division septa (58, 69–73). By binding to an ephemeral signal (UndP, the lipid carrier that anchors sugar moieties of peptidoglycan precursors) our findings suggest that ShfA localization is closely coupled to active peptidoglycan synthesis rather than being strictly restricted to a particular subcellular region. Consistent with this notion, when the coordination between peptidoglycan synthesis and membrane invagination was disrupted (either genetically or by addition of antibiotic) ShfA failed to localize properly. Moreover, the observation that ShfA can be artificially deployed to aberrant sites of lateral cell wall incorporation is further consistent with the idea that ShfA localization is guided by sites of active cell wall synthesis rather than a static cellular landmark. A parallel to this scenario has been reported for the polar or septal localization of the PknB kinase in *Mycobacterium tuberculosis* and *Staphylococcus aureus*, where PknB binds to the lipid II cell wall precursor via its extracellular PASTA domain (74, 75). However, although lipid II is anchored to the membrane via undecaprenyl phosphate, the interaction of PknB with lipid II occurs through the extracellular muropeptide portion of lipid II, rather than through the hydrophobic transmembrane lipid portion. Another similar situation exists with the reported preferential interaction between the lipid cardiolipin and pole-localized proteins ProP and MinD in *E. coli* (76, 77). The conical shape of this lipid has been invoked in its preferential accumulation at negatively curved regions of the cell (78). However, the ability of other anionic lipids such as phosphatidylglycerol to compensate for the absence of cardiolipin in a cell to interact with proteins and drive their proper localization and the detection method for cardiolipin in vivo have complicated understanding if the precise role of cardiolipin in the localization of these proteins is direct or indirect (79).

The ability of ShfA to spontaneously insert into the membrane near sites of active peptidoglycan synthesis in multiple organisms, combined with the universal conservation of UndP in bacteria raises the possibility of exploiting this activity to develop bacterial cell biological methods that can function in different species. For example, ectopic production of ShfA-GFP may reveal sites of active peptidoglycan assembly in real time; combining the ectopic production of ShfA with, for example, proximity ligation (80) may enable the identification of proteins near the site of active peptidoglycan synthesis in diverse bacterial species. Another intriguing possibility is the development of a fluorescence microscopy-based two-hybrid assay to screen for protein-protein interactions. Unlike previously reported systems that require the screening to be performed in rod-shaped cells or in a model bacterial species like *E. coli* (81), the ectopic production of ShfA fused to a bait protein may be used to recruit interacting proteins in preassembled libraries of gene fusions to fluorescent reporters to the division septum, which may be visualized by high throughput fluorescence microscopy in the native environment of the bacterium being studied.

An outstanding question is the precise nature of the “antidote” function of ShfA during sporulation. Previously, we reported that the ShfP phosphoesterase is responsible for the extracellular release of millimolar quantities of glycerol that functions as a signaling molecule to delay sporulation in other cells that had not yet committed to the sporulation program (32). This glycerol-mediated inhibition occurs via the KinD histidine kinase. However, the very cells that produce extracellular glycerol were also susceptible to an apparent toxicity of the glycerol during sporulation, which requires ShfA to counteract. Interestingly, ShfP, a cell surface calcineurin-like phosphoesterase distributed widely across most major bacterial lineages, is not absolutely conserved among sporulating Bacillota (32). In contrast, ShfA is broadly conserved among sporulating Bacillota (82), suggesting that it performs an essential function during sporulation that is independent of the glycerol antidote function that we previously described (32). One possibility for ShfA function arises from our previous observation that, during sporulation, the nanoenvironment immediately surrounding the forespore surface is more acidic than the rest of the mother cell cytoplasm (83), which we reported was unfavorable to the peptidoglycan biogenesis reactions that occur at that location that are required for cortex assembly. As a result, the sporulation-specific AAA+ chaperone SpoVK is required to activate the glycosyltransferase MurG in this nanoenvironment for successful lipid II synthesis during cortex assembly. Previous observations have indicated that the phosphate head group of UndP is sensitive to even mild acidification (84, 85). Thus, the local acidification around the forespore likely destabilizes UndP (and likely also the UndP-containing peptidoglycan precursors lipid I and lipid II) (Fig. 7). We therefore hypothesize that ShfA may function as a membrane-embedded factor that buffers the local pH near the outer forespore membrane and/or physically shields UndP from the acidic nanoenvironment at the forespore surface to stabilize UndP and promote the lipid I and lipid II synthesis activities of MraY and MurG, respectively. Notably, although poorly conserved, the variable cytoplasmic tails of ShfA orthologs have a net mean positive charge of 10.5. Indeed, this excess positive charge might help locally neutralize the acidic nanoenvironment, thereby allowing UndP stabilization. Hence, the identification of UndP as the localization signal for ShfA might guide future work in determining a precise broadly conserved molecular function for ShfA during sporulation that involves assembly of the cortex cell wall and raises the possibility that UndP might be a widespread localization cue for membrane proteins across bacteria.

**Figure 7.**
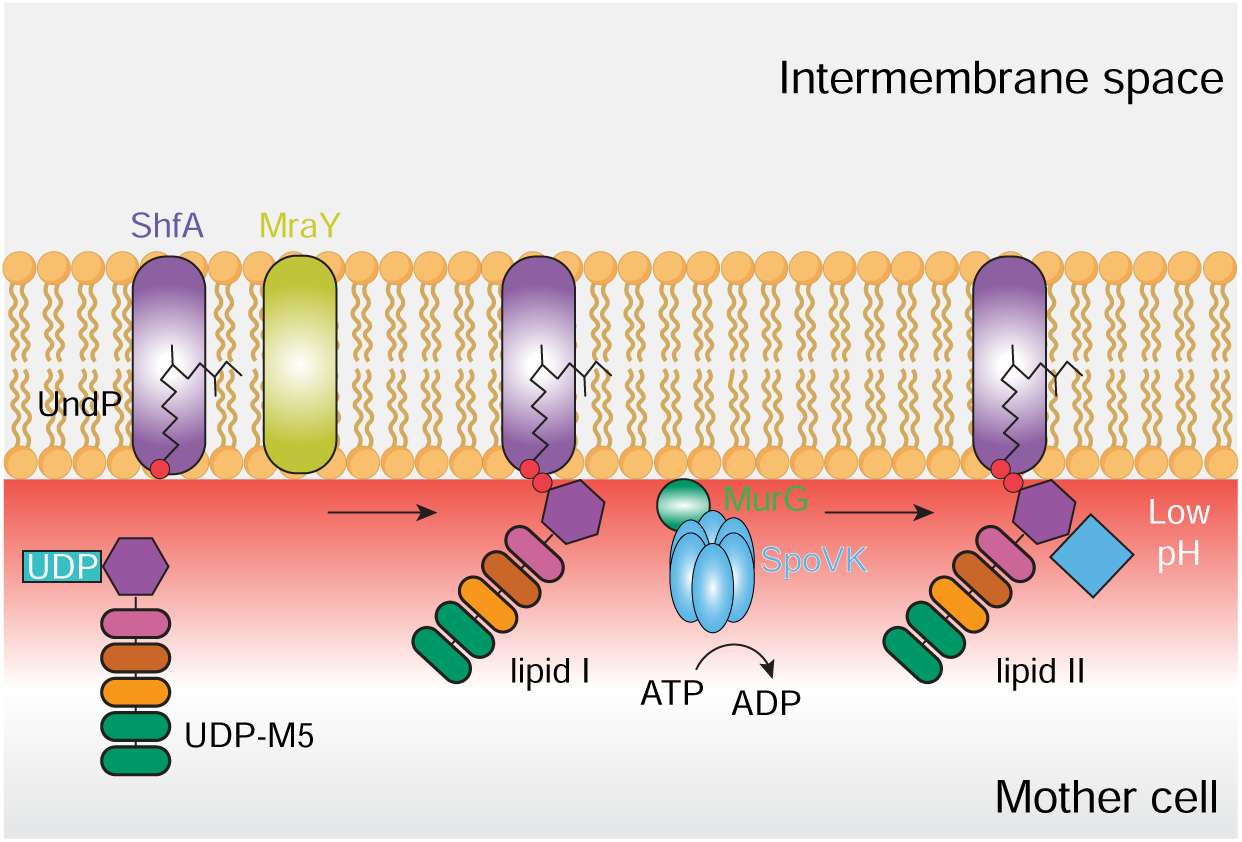
Model of UndP-mediated recruitment of ShfA to the outer forespore membrane during sporulation in *B. subtilis*. ShfA (purple) is targeted to the outer forespore membrane where it likely directly binds the lipid carrier UndP and stabilizes UndP in the low pH mother cell cytosolic nanoenvironment surrounding the forespore. UndP is then employed as a precursor by the integral membrane protein MraY (light green) to generate lipid I; lipid I is then converted to lipid II by the membrane-associated MurG, which requires its sporulation-specific AAA+ chaperone SpoVK for activation in the acidic nanoenvironment.

## MATERIALS AND METHODS

### Strain construction and general methods

Strains used in this study (genotypes listed in Table S1) are otherwise isogenic derivates of *B.subtilis* PY79 strain (86). Genes of interest were PCR amplified to include with either their native or heterologous promoter and cloned using Gibson Assembly kit (NEB) into integration vectors: pDG1731 (for insertion at *thrC* locus), pDG1662 (for insertion into the amyE locus), pDR111 (for insertion at *amyE* locus, downstream of IPTG inducible *P_hyperspank_* promoter) (87–89). Site directed mutagenesis was performed using the QuikChange kit (Agilent). All plasmids were integrated into *B. subtilis* (PY79) chromosome by double recombination event at the specified ectopic locus. For determination of membrane topology of ShfA, various C terminal truncations were PCR amplified and cloned in-frame at the N-terminus of the dual reporter PhoA–LacZα vector (pkTop), between *Hind*III and *Bam*HI sites. To overexpress *B. subtilis shfA*-*gfp* in *S. aureus*, *shfA-gfp* was PCR amplified and inserted into cadmium-inducible plasmid pJB67 (90). Plasmid was introduced into *S. aureus* RN4220 by electroporation. To overexpress *B. subtilis shfA-gfp in E. coli*, *shfA-gfp* was PCR amplified from *B. subtilis* strain CW205 and inserted into the IPTG inducible pET28A vector. The resulting clone was further transformed into *E. coli* overexpression strain BL21(DE3) (New England Biolabs). All plasmid constructions were verified by DNA sequencing before chromosomal integration or transformation.

### Growth conditions

*B. subtilis* PY79 or isogenic mutant derivates were grown on Lysogeny Broth (LB) agar plates (10 g tryptone, 10 g yeast extract, 10 g NaCl per liter; KD Medical) for single colony isolation. Single colonies were inoculated in 2 ml casein hydrolysate medium (CH; KD Medical) and grown overnight at 22 °C rotating at 250 rpm. Overnight cultures were diluted 1:40 in 10 ml of CH medium (to OD_600_ = 0.05) and grown for 2 h at 37 °C or 25 °C (for *divIC* temperature sensitive construct). For *E. coli* (strain BL21(DE3)) and *S. aureus* (strain RN4220), overnight cultures were grown in LB broth and tryptic soy broth (TSB; KD Medicals) and diluted 1:40 in the respective medium and grown to OD_600_ = 0.4.

For the targeted insertion experiment during sporulation, the strains were induced with 1 mM IPTG or 0.5% xylose at time 3.5 h post sporulation and imaged 30 min post induction. For the undecaprenyl phosphate (UndP) depletion experiment, 0.1 mM IPTG was supplemented throughout the experiment and removed at 2 h post sporulation induction for UndP depletion, cells were imaged 30 min post IPTG removal.

For localization experiments during vegetative growth in *B.subtilis*, *E. coli* and *S. aureus*, *shfA* expression was induced with 0.5 mM IPTG, 1 mM IPTG and 1 μM of cadmium chloride respectively. For antibiotic related localization experiments, 0.5 μg ml^-1^ vancomycin and 16 μg ml^-1^ of tunicamycin were added along with IPTG and imaged 30 min post induction. ShfA localization in a *divIC* temperature sensitive background, post induction with IPTG, cultures were shifted to 37 °C for 1 h before imaging. For MreB experiment, cells were imaged after 2 h post induction. For single molecule experiments during sporulation, the cells were harvested after 3 h post sporulation induction.

### Membrane topology determination

*E.coli* DH5α F’Iq (NEB) strain was transformed with the pKTop vector carrying the relevant ShfA-PhoA–LacZα fusion constructs and plated on LB agar plates containing 50 μg ml^-1^ kanamycin and 0.2% glucose and incubated for 18-20 h at 30 °C. 2-3 colonies were inoculated in 2 ml Lennox broth containing kanamycin (50 μg ml^-1^) and glucose (0.2 %) for 5-6 h at 30 °C. The culture was further diluted to OD_600_ = 0.05. 5 μl of diluted culture was spotted on a fresh dual indicator plate, containing kanamycin (50 μg/mL), IPTG (1 mM), BCIP (80 μg ml^-1^, Sigma-Aldrich) and Red-Gal (100 μg ml^-1^, Sigma-Aldrich), incubated for 18-20 h at 30 °C.

### Sporulation efficiency assay

*B. subtilis* cells were induced to undergo sporulation in Difco Sporulation Medium (DSM; K.D. Medical) for at least 24 h. Non-sporulating and sporulation-defective cells were killed by incubating the culture at 80 °C for 20 min. Cultures were then serially diluted and colony forming units (CFU) were enumerated to calculate viable spore count. Sporulation efficiency was determined as a ratio relative to the CFU obtained in a parallel culture of the WT strain (PY79).

### Epifluorescence microscopy

Cells were induced to sporulate using the resuspension method (91) in Sterlini-Mandelstam (SM) medium (KD Medical). Cells grown in casein hydrolysate (CH) medium were harvested by centrifugation at 4000 × *g* and pellets were resuspended in 10 ml SM supplemented with threonine (80 µg ml^-1^; Sigma – Aldrich), as required if the strains harbored an insertion that disrupted the *thrC* locus, and grown for 2.5-3 h. 500 μl of the cultures were centrifuged at 4000 × *g*. Cell pellets were resuspended in 100 μl 1 × PBS and stained with 2 μl of 10 mM HADA to visualize site of active peptidoglycan incorporation for 5 min at 37 °C. HADA stained cells were harvested by centrifugation and washed twice with 0.5 ml 1 × PBS. The pellet was resuspended in 100 μl 1 × PBS containing fluorescent dye FM4-64 (1 µg ml^-1^; Invitrogen) and/or MitoTracker Deep Red FM (1 µg ml^-1^; Invitrogen) respectively. 3 μl was spotted on a poly-L-lysine coated glass bottom dish (MaTek Corp.) and covered with a 1% agarose pad (made with distilled water) and imaged at 25 °C. The cells were viewed with a DeltaVisison Core microscope system (Applied Precision) as described previously (92). Ten planes were acquired every 0.2 μm. The data was deconvolved using SoftWorx software (GE Healthcare). Data analyses (septal localization quantifications) were performed using Fiji software (NIH).

### Single molecule fluorescence microscopy

Single molecule microscopy was performed largely as described previously (29). Single-molecule localization imaging was carried out on a custom Nikon Ti-E microscope equipped with an N-STORM module and a CFI Apo TIRF 100× 1.49 NA oil-immersion objective. Samples were illuminated in highly inclined mode with a 561 nm laser (Coherent, USA). Images were recorded on a scientific CMOS camera (Prime 95B, Teledyne Photometrics) with a 20 ms exposure time, collecting 400 frames per acquisition. A 637-nm laser (Coherent, USA) was used as the excitation wavelength for imaging the far-red channel for membrane staining. For live cell imaging of sporulating cells 3 h post-induction, 300 µl of sporulating culture was harvested by centrifugation. For HaloTag-conjugated protein visualization, cells were resuspended in 30 µl of 200 pM JFX554 (Promega). Simultaneously, 30 µl of 500 nM of MitoTracker Deep Red FM (ThermoFisher Scientific) was added to stain membranes. The samples were incubated at 37 °C for 15 min. Labeled cells were washed three times with 1× PBS and resuspended in 50 µl 1× PBS. 3 µl of cells were pipetted onto a plasma cleaned coverslip and covered with a 1% (w/v) agarose pad to immobilize the cells. Data were analyzed as previously described (29). Briefly, single molecules were first localized with 2D Gaussian fitting subject to a log-likelihood ratio test with a ‘localization error’. A maximal expected diffusion constant was set to connect localizations between consecutive frames. Regions of interest (around forespores and mother cells) were generated from far red membrane staining images. These regions were segmented and used as a mask to filter tracking data for individual single molecule data. Tracks were classified into three different categories: mobile, forespore; mother cell; and immobile, forespore.

### ELISA assay

GFP ELISA Kit (Cell Biolabs, Inc.) was used to quantify the protein levels of wild type ShfA-GFP and ShfA-GFP variants according to manufacturer’s protocol. Absorbance at 450 nm (Synergy H1 microplate reader, BioTek) reading was taken for the reactions and the GFP standards. Slope estimated from the GFP standard curve was used to calculate the GFP levels of the strains.

### Computational Analysis

#### Sequence Analysis and Homology Detection

Sequence profile searches were performed against the NCBI non-redundant (nr) protein database (93), and the nr50 clustered database (nr clustered with MMseqs2 at 50% sequence identity), and a curated database containing 4210 complete prokaryotic genomes, 230 select eukaryotic genomes and 2983 curated metagenomes using PSIBLAST (94) and hidden Markov model (HMM) searches via JACKHMMER (95). To manage dataset size and define relationships, we employed the MMseqs program (96) for clustering by bitscore or percentage similarity. Redundancy was removed using a 70% identity and 80% coverage threshold, while homologous groups were established with an e-value of 10^−^³ and 80% coverage. Multiple sequence alignments were constructed using either FAMSA (97) or MAFFT (98) (local-pair algorithm with –maxiterate 3000, –op 1.5, and –ep 0.2).

#### Domain Identification and Comparative Genomics

Known domains were identified using Pfam HMMs (99) and a curated internal collection of HMMs and PSSMs (Aravind laboratory). These were searched using RPSBLAST (100). Genomic neighborhood extraction and analysis were automated via the ROTIFER and TASS packages. Gene neighborhoods were clustered using MMseqs2 with a maximum intergenic distance of 150 nucleotides, followed by ORF orientation filtering. Phylogenetic analysis was performed using FastTree (101) and iqTREE2 (102).

#### Structural Modeling

3D structural modeling was executed with AlphaFold3 (103). Model reliability was rigorously evaluated using pLDDT (local chain confidence), PAE (domain orientation), and ipTM/PDI metrics for lipid-protein interface confidence. Structural visualization and analysis were performed in MOL* (104). Prediction of transmembrane helices and membrane topology was performed using the Phobius and DeepTMHMM programs.

## Data, Materials, and Software Availability

This work does not report original code. All other data are included in the manuscript and/or supporting information.

## ACKNOWLEDGEMENTS

We thank members of the KSR lab for suggestions and comments on the manuscript, S. Gottesman and A. Khare for discussions, and D. Rudner and C. Weiss for strains. This research was supported by the Intramural Research Program of the National Institutes of Health (NIH), National Cancer Institute, Center for Cancer Research (K.S.R.) the National Library of Medicine (L.A.), and the National Institute of Biomedical Imaging and Bioengineering (J.C.). The contributions of the NIH author(s) are considered Works of the United States Government. The findings and conclusions presented in this paper are those of the authors and do not necessarily reflect the views of the NIH or the U.S. Department of Health and Human Services.

## Supporting Information

**Figure S1.**
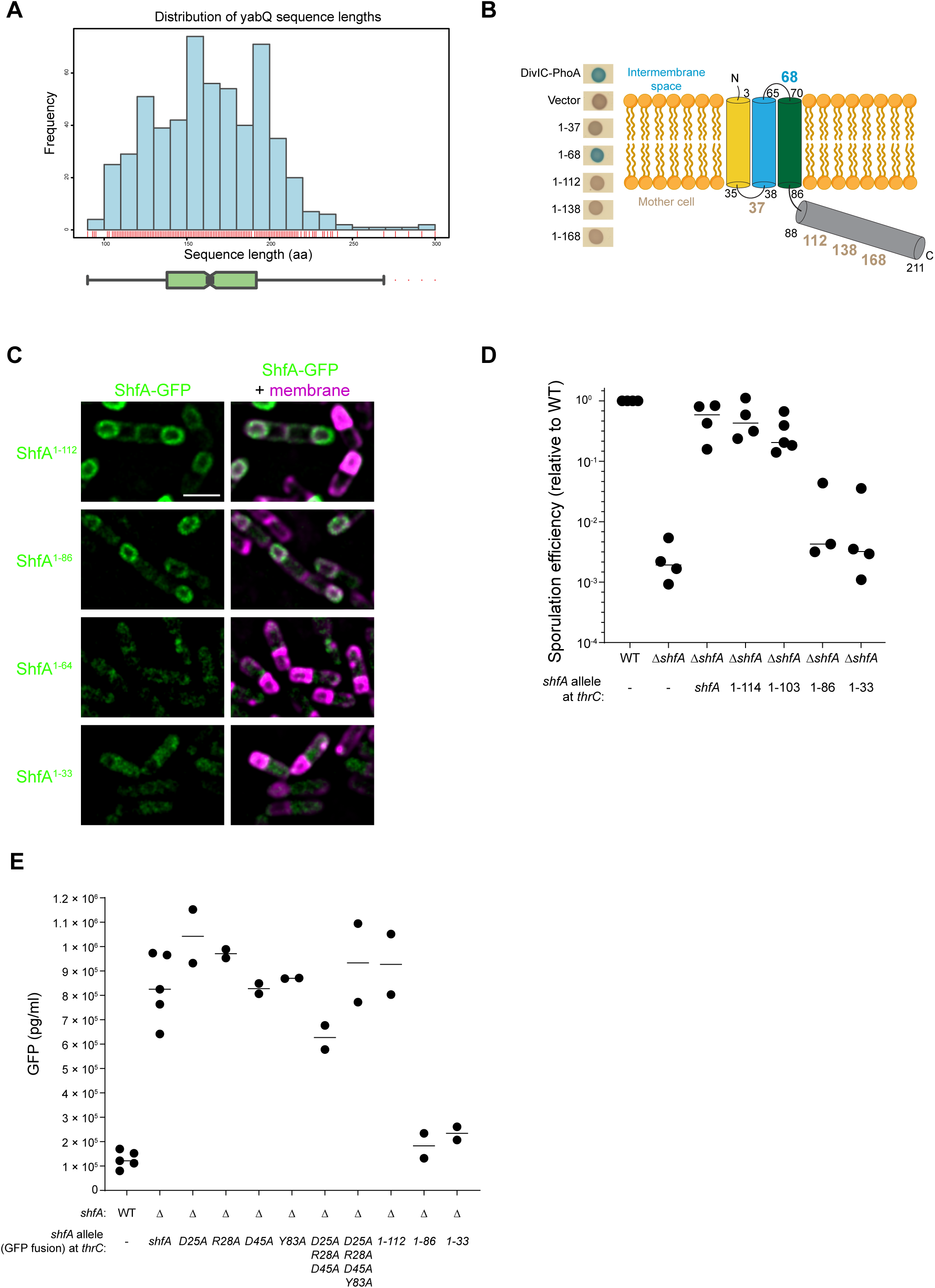
The YabQ domain of ShfA is important for function and subcellular localization. (A) Histogram of ShfA protein lengths in amino acids from a curated, nonredundant set of complete genomes illustrates variability across species. A rug plot along the X-axis (red) indicates the density of observations in each bin, and a green box plot below shows the length distribution with red dots marking outliers. (B) Left: representative *E. coli* colonies harboring indicated fusions of ShfA to PhoA and plated on agar containing 5-bromo-4-chloro-3-indolyl phosphate to indicate PhoA activity (blue color indicates high PhoA activity). Right: Schematic of the inferred ShfA membrane topology. Amino acid positions are indicated; fusions to ShfA residues indicated in brown resulted in low PhoA activity, suggesting cytosolic localization of PhoA; fusion to ShfA residue indicated in blue resulted in high PhoA activity, suggesting extracytosolic localization of PhoA. C-terminal fusion of PhoA to DivIC is a positive control for extracytosolic localization of PhoA; “Vector” is a negative control for cytosolic localization of PhoA. Strains: VP254, VP273, VP274, VP275, VP276, VP277 and VP278.(C) Subcellular localization of indicated ShfA-GFP truncations 3.5 h after induction of sporulation. Left: fluorescence from ShfA-GFP; right: overlay, GFP and membranes. Strains: CW214, CW212, CW213, CW211. (D) Sporulation efficiencies relative to WT for strains harboring the indicated truncation allele of *shfA*. *thrC* is a chromosomal locus in which different alleles of *shfA* are inserted. Bars represent mean; data points indicate independent experiments. (E) Intracellular accumulation of indicated ShfA-GFP variant produced from the *thrC* locus at 3.5 after induction of sporulation as determined by ELISA.

**Figure S2.**
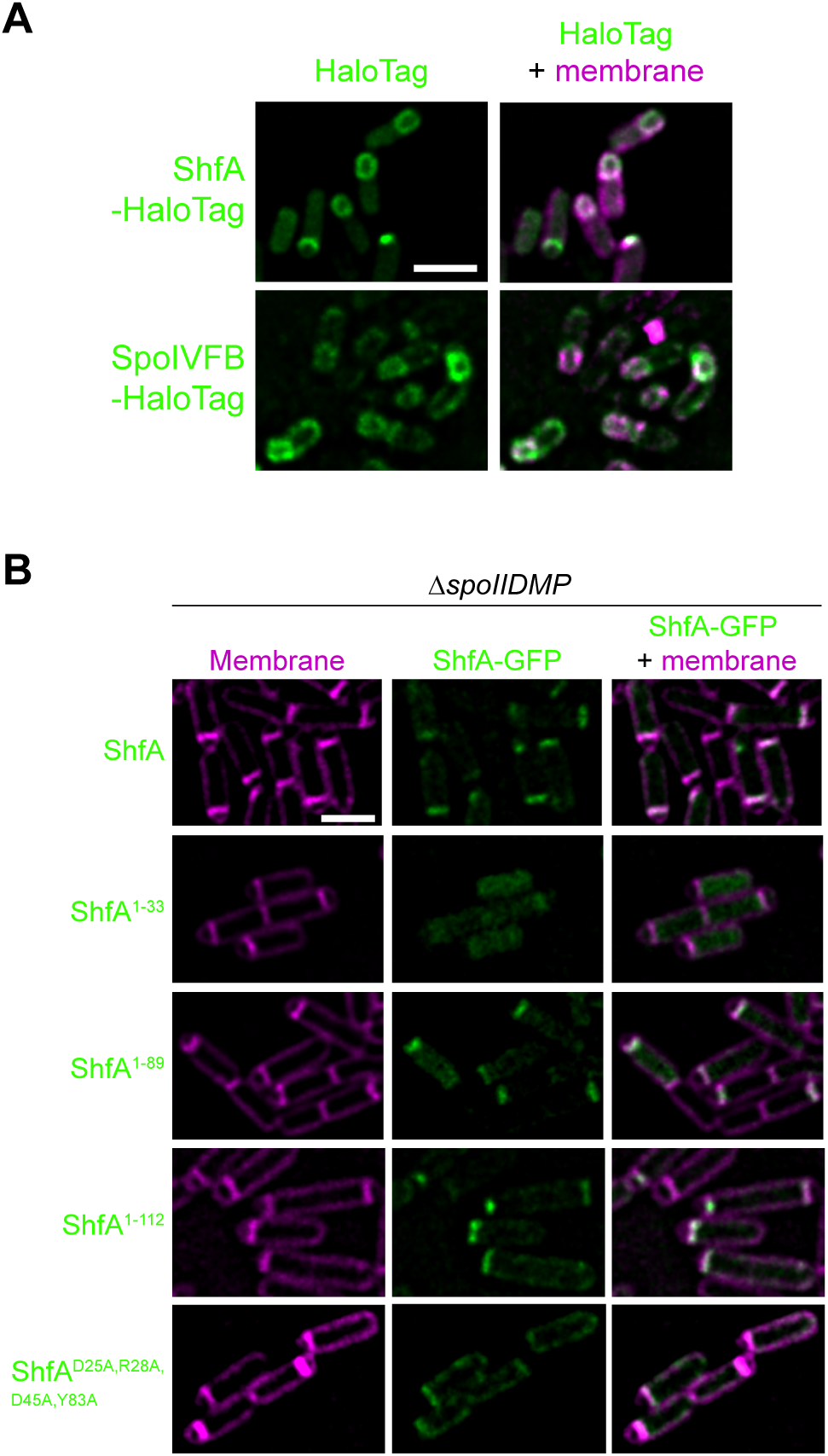
ShfA-GFP can localize to a flat asymmetric septum during sporulation. (A) Subcellular localization of ShfA-HaloTag and SpoIVFB-HaloTag in sporulating *B. subtilis* strains 3.5 h after induction of sporulation in the presence of excess JFX554 dye (100 nM). Left: fluorescence from JFX554 dye bound HaloTag; right: overlay, HaloTag and membranes visualized with MitoTracker Deep Red FM. Strains: VP461 and VP462. Scale bar: 2 µm. (B) Subcellular localization of indicated ShfA-GFP truncations or variant 3.5 h after induction of sporulation in a Δ*spoIID/M/P* strain. Left: membranes visualized using FM4-64; center: fluorescence from ShfA-GFP; right: overlay, GFP and membranes. Scale bar: 2 µm.

**Figure S3.**
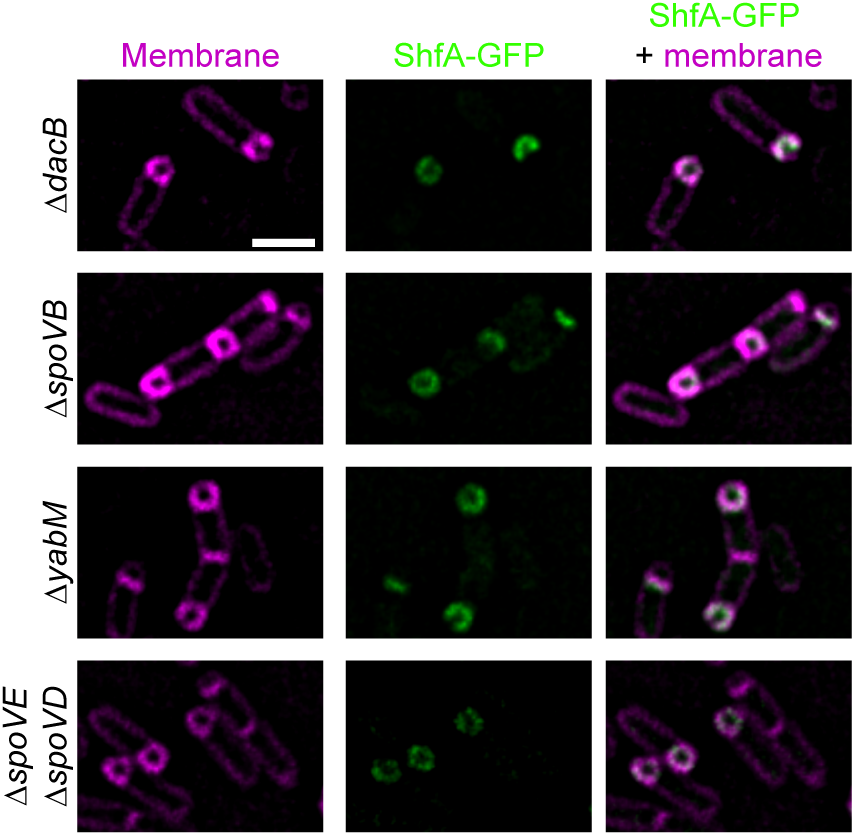
Spore cortex peptidoglycan remodeling proteins do not determine ShfA localization. Subcellular localization of ShfA-GFP 2.5 h after induction of sporulation in the absence of *dacB*, *spoVB*, *yabM*, or in a Δ*spoVE* Δ*spoVD* double mutant. Left: membranes visualized using FM4-64; center: fluorescence from ShfA-GFP; right: overlay, GFP and membranes. Strains: VP118, VP131, VP123, VP137. Scale bar: 2 µm.

**Table S1.**
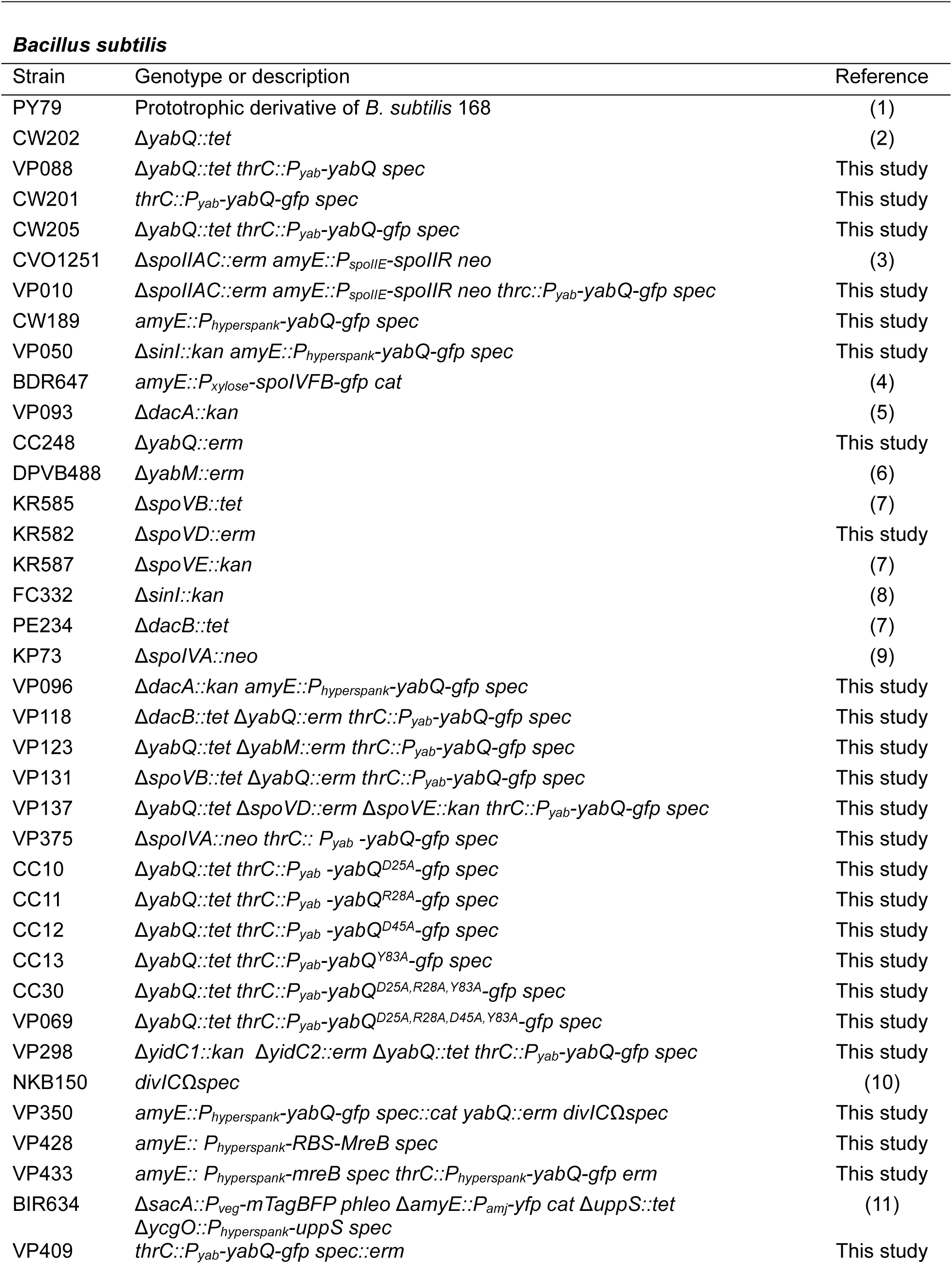
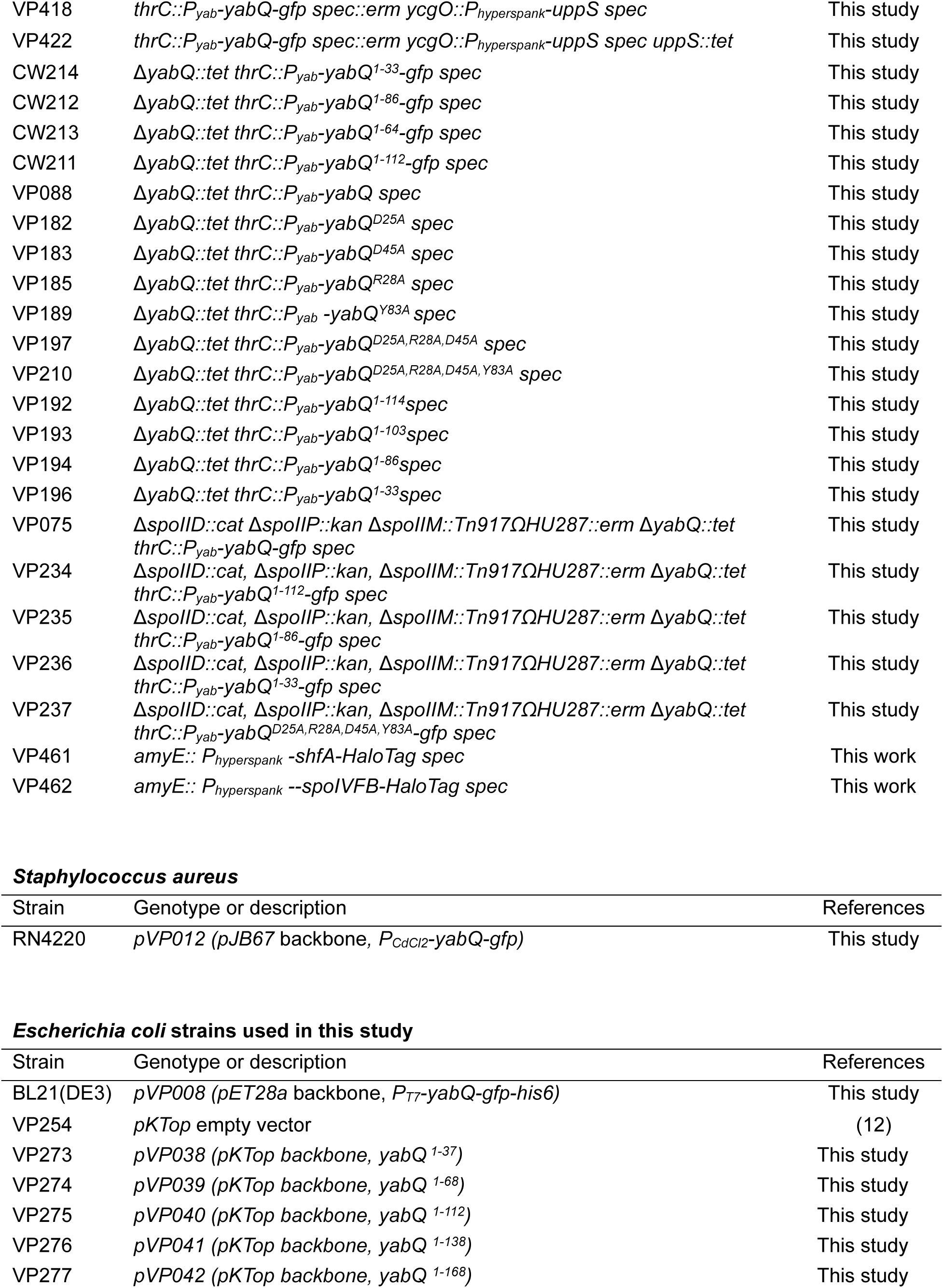
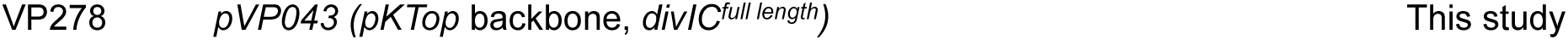
Bacterial strains used in this study.

